# Image-based assessment of plant disease progression identifies new genetic loci for resistance

**DOI:** 10.1101/2021.07.13.452064

**Authors:** Valérian Méline, Denise L. Caldwell, Bong-Suk Kim, Sriram Baireddy, Changye Yang, Erin E. Sparks, Edward J. Delp, Anjali S. Iyer-Pascuzzi

## Abstract

A major challenge in global crop production is mitigating yield loss due to plant diseases. One of the best means of disease control is plant resistance, but the identification of genes that promote resistance has been limited by the subjective quantification of disease, which is typically scored by the human eye. We hypothesized that image-based, non-destructive quantification of disease phenotypes would enable the rapid identification of new disease resistance loci. We tested this using the interaction between tomato and *Ralstonia solanacearum*, a soilborne pathogen that causes bacterial wilt disease. We acquired over 40,000 time-series images of disease progression in a tomato recombinant inbred line population, and developed an image analysis pipeline providing a suite of ten traits to quantify wilt disease based on plant shape and size. Quantitative trait loci (QTL) analyses using image-based phenotyping identified QTL that were both unique and shared compared with those identified by human assessment of wilting. When shared loci were identified, image-based phenotyping could detect some QTL several days earlier than human assessment. Thus, expanding the phenotypic space of disease with image-based, non-destructive phenotyping allowed both earlier detection and identified new genetic components of resistance.

## Introduction

Plant diseases are a significant global constraint to crop production. Developing disease resistant crops requires identifying the plant genomic regions and genes that contribute to resistance to the pathogenic microbes that cause disease. This in turn depends on phenotyping large populations of plants for their responses to pathogens. Phenotyping plant diseases is challenging because diseases cause complex, quantitative phenotypes that can occur at different scales – e.g. on parts of leaves, entire leaves, or the whole plant. In addition, disease phenotypes vary over time and depend on environmental conditions, plant age, and pathogen virulence. Disease symptoms such as wilting or necrotic spots have traditionally been scored with the human eye, but these scores are subjective, can vary by individual, and are difficulty to accurately quantify.

The challenging nature of visual disease assessment has led to the use of sensors including RGB, hyperspectral, chlorophyll fluorescence and thermal cameras to assess disease symptoms (Colwell, 1956; Jackson, 1986; Bock et al., 2010; Simko et al., 2017). Compared to assessment by the human eye, image-based phenotyping is faster, more reproducible, and more sensitive to small variations in disease symptoms that can be critical for detecting resistance loci (Bock et al., 2008, 2010; Stewart and McDonald, 2014; Stewart et al., 2016; Simko et al., 2017; Shakoor et al., 2017). Many studies have used or developed tools to assess plant symptoms using different types of sensors (Mahlein, 2016; Mahlein et al., 2017, 2019; Lowe et al., 2017; Shakoor et al., 2017; Mochida et al., 2019; Mir et al., 2019; Pérez-Bueno et al., 2019; Pineda et al., 2021; Simko et al., 2017). However, few studies have used these technologies in QTL or Genome Wide Association (GWA) analyses for responses to plant pathogens, and all have used destructive methods (Yates et al., 2019; Fordyce et al., 2018; Corwin et al., 2016). It has remained challenging to use image-based, non-destructive phenotyping for disease resistance across large populations, both because of technical factors like the expense of phenotyping platforms and the time associated with imaging, and also biological factors such as differences in plant morphology and disease progression within a population.

The soil-borne betaproteobacterium *Ralstonia solanacearum* is the causal agent of bacterial wilt disease and has been ranked as one of the top 10 most destructive plant bacterial pathogens of all time (Mansfield et al., 2012). *Ralstonia* infection causes susceptible plants to wilt, and the amount of wilting correlates with a plant’s level of susceptibility (Genin, 2010; Genin and Denny, 2011). The bacterium is a major production constraint in Solanaceous crops both globally and in the United States, where disease loss in tomatoes can exceed 70%. In crops, resistance to *Ralstonia* is quantitative, but the quantitative trait loci (QTL) underlying resistance to US strains of *Ralstonia* are largely unknown. QTL for other strains have been mapped (Danesh et al., 1994; Thoquet et al., 1996a, 1996b; Mangin et al., 1999; Wang et al., 2000; Carmeille et al., 2006; Jaw-Fen Wang et al., 2013; Shin et al., 2020) but have not been cloned, and the host determinants necessary for resistance remain mostly unspecified.

The limited identification of QTL for *Ralstonia* resistance can be attributed in part to the difficulty in accurately scoring plant wilting. Wilting is traditionally measured on a 0 – 4 scale, in which 0 indicates a plant with no wilting, 1 = 1 – 25%, 2 = 26 – 50%, 3 = 51 – 75% and a score of 4 indicates a plant with 76 – 100% wilted leaves (Schandry, 2017). While it is straightforward to assess the ends of the spectrum, rating plants with scores of 2 or 3 is particularly difficult. This is due to the subjective nature of visually determining when a leaf has lost sufficient turgor to qualify as wilted. Reliable disease phenotyping is critical for identifying QTL for resistance to *Ralstonia* and the development of resistant varieties.

Here, we used image-based, rapid, non-destructive phenotyping to identify new tomato genetic resistance loci to *Ralstonia*. We developed a rapid, semi-automated imaging and trait analysis pipeline to quantify bacterial wilt disease in a recombinant inbred line (RIL) population derived from *Ralstonia*-resistant and susceptible tomato genotypes. We found both unique and shared QTL between our image-based traits and plant wilting scored by the human eye. At least one of the QTL was detected by image-based phenotyping before the onset of visual symptoms, demonstrating that image-based phenotyping captures the disease phenotype at early stages of infection. These results demonstrate that imaged-based, non-destructive phenotyping can shed light on new aspects of disease and improve our ability to identify genetic loci crop resistance.

## Results

### Development of an aboveground imaging and semi-automated analysis pipeline

We first constructed a simple, low-cost imaging system that allowed us to semi-automate aboveground disease phenotyping. Each plant was placed on a commercially available turntable, and plants were imaged with a Canon DSLR (Supplemental Figure 1; details in methods). The turntable and camera were connected with Photocapture 360 (Ortery technologies), which allowed us to automatically capture images every 45 degrees (8 images per plant). Using this system, we were able to non-destructively image each plant in less than 2 minutes, with minimal manual labor. Each image included a fiducial marker for post-image color correction. Plants in the F9 generation from a RIL population derived from a cross between resistant Hawaii 7996 (H7996) and susceptible West Virginia 700 (WV) were imaged the day before inoculation with *R. solanacearum* strain K60, and at 3, 4, 5 and 6 days post inoculation (dpi). At 3 dpi, symptoms were not present in susceptible parent WV, but by 6 dpi these plants were completely wilted. We imaged five replicate plants of each RIL as well as the parental lines. Using this system, we captured over 40,000 images for high-resolution disease phenotyping. The same set of plants was also visually scored by the human eye at 8 dpi. Visual scoring was based on the percentage of wilted leaves. Tomato phenotypes in the RIL population ranged from highly susceptible to highly resistant, consistent with the quantitative nature of disease resistance (Supplemental Figures 2 and 3).

We next developed a set of mathematical descriptors to phenotype wilting over time in our images. Plant wilting is a composite phenotype, and we used 10 image-based traits (Supplemental Table 1) to describe different aspects of wilting: convex area, convex width, plant area, plant height, plant width, X mass, Y mass, center of mass (CM) height, CM width, and color. Several of these, such as the area and width of the convex hull, are traditional methods of describing aboveground plant shape. Because the center of mass of a plant leaf changes as a plant wilts, we developed additional descriptors based on the distance of the leaf center of mass from the stem (CM width and CM height, X mass and Y mass). We then developed a pipeline which used the original image as input, performed color correction, and quantified each descriptor (Figure 1 and Supplemental Figure 4, and methods).

**Figure 1:**
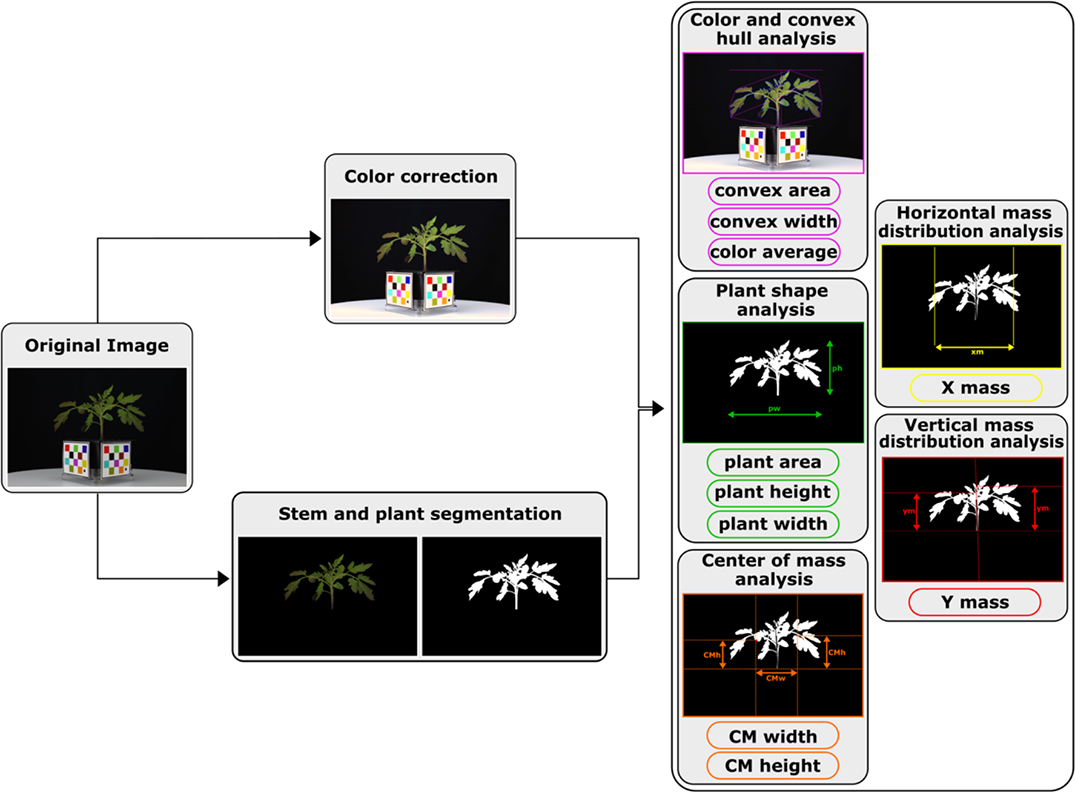
Diagram of the semi-automated analysis pipeline and 10 image descriptors

### Image-based traits differentiate resistant and susceptible plants

To validate the efficiency of the image-based descriptors to estimate wilting phenotypes, we tested whether image-based phenotyping descriptors could differentiate resistant from susceptible plants. Supplemental Figure 5 shows the average normalized score for each of the image-based descriptors for each parent and the RIL population from -1 (the day before inoculation) to 6 dpi. Most descriptors, particularly those based on plant width or convex hull, had clearly divergent values in resistant and susceptible plants at a given time point. RIL descriptor values ranged from those of the resistant to susceptible parents and occasionally showed transgressive segregation (Supplemental Figure 5). Because plant shape at 6 dpi depends on shape of the same plant at -1 dpi, we used the evolution of each descriptor from day -1 to 6 dpi in our QTL analysis. This evolution was termed a ‘trait’. Trait values were clearly different for resistant and susceptible parents for all traits except color (Figure 2), which was not used further.

**Figure 2:**
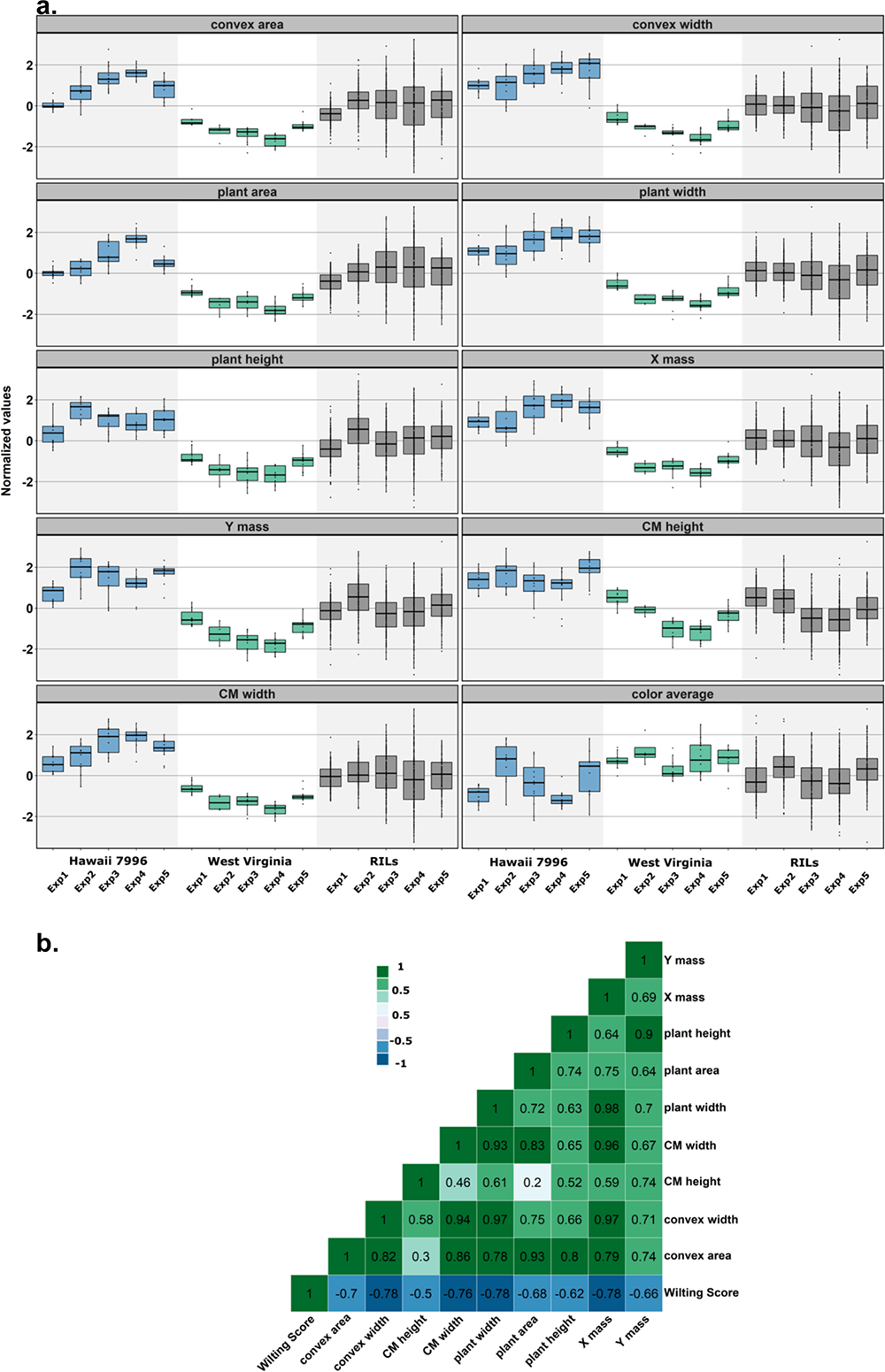
Trait evolution during disease and correlation among descriptors. a. Boxplots showing the evolution of the normalized values between -1 and 6 dpi for each of the five biological replicates for the ten image-based traits for resistant Hawaii 7996, susceptible West Virginia parents and the 166 RILs. Color was unable to differentiate resistant from susceptible plants and was not used further. b. Heatmap showing the Pearson correlation values between image-based traits used in our QTL analysis and the human eye based wilting score in 2019. The correlations values were determined using the same plants imaged and visually assessed in 2019.

### Image-based phenotyping reveals drivers of the wilting phenotype

We next investigated whether any of our traits were major components of the wilting phenotype. We visually scored plants and asked how well our image-based traits correlated with human visual scoring. Wilting is categorized by loss of plant leaf turgor that results in drooping leaves, and decreased plant width and height. Determining how much a plant has wilted is challenging, in part because it can be difficult to quantify how much each a leaf has drooped and how much drooping of one leaf correlates with whole plant wilting.

We aimed to quantify leaf drooping using center of mass traits. Among our image-based traits, those which were functions of the leaf center of mass were highly inversely correlated with visual wilting (i.e. as a plant wilts, the center of mass decreases), suggesting that these are major drivers of the wilting phenotype. These traits included CM width, plant width and X mass (r > - 0.75) (Figure 2B).

Several of our traits describe similar aspects of plant shape, such as height or width, through different methods. These traits tended to be highly correlated with each other. For example, plant height vs Y mass use different methods to describe plant width (based on the plant mask or the center of mass of the stem masks; see Methods), and were highly positively correlated with each other (0.93; Figure 2B).

We trained a random forest, consisting of 1000 decision trees, to use data from 6 dpi to predict the expert visual score assigned at 8 dpi. We used 969 plants in a 60:40 training-testing data split, and achieved a classification accuracy of 83%. To identify the traits that provided the most efficient estimation of wilting symptoms, we randomly scrambled the nine image-based traits one at a time. Plant width and plant area had the most impact on wilting score prediction (Supplemental Figure 6).

### QTL analysis identifies 20 wilting QTL in 10 clusters across the tomato genome at 6 dpi

Our overall goal was to identify tomato genomic regions that provide resistance or susceptibility to *Ralstonia*. Prior to this analysis, we first generated a genetic map using Genotyping-by-Sequencing (GBS). We identified 632 high-quality SNPs for linkage mapping using GBS. We combined these with 112 SolCap markers, and subsequently generated a linkage map using ICI mapping software (Meng et al., 2015). Our linkage map consisted of ∼1300 cM (Supplemental Figure 7) with an average per chromosome marker density that varies from 1.8 to 7.48 cM (Supplemental Table 2).

For QTL analysis, we mapped the evolution of each our nine image-based descriptors from -1 to 6 dpi. In addition to the image-based phenotyping, we used two years (2016 and 2019) of visually scored phenotyping data at 8 dpi. Visually assessed wilt scores from 2019 were quantified from the same plants that were used for image-based phenotyping. In 2016, plants were only visually scored for wilting, and no image-based measurements were taken. Using ICI mapping software and composite interval mapping (CIM) for all traits, we identified 20 QTL within the RIL population with a LOD score above 3 (Table 1). To be consistent with previous studies of tomato-*Ralstonia* QTL mapping, we call these QTL, ‘Bacterial wilt resistance (*Bwr*) QTL’. Each *Bwr* QTL explained approximately 6 to 11% of the variation in response to *R. solanacearum* strain K60, and together the clusters explained more than 88% of the variation (calculated using the sum of the QTL with the highest PVE in each cluster).

**Table 1.** Overview of the 20 wilting QTL identified in 10 clusters across the tomato genome at 6 dpi. LOD: maximum value of the Logarithm of the odd. PVE: Percentage of phenotypic variance explained. Add: Additive effect. Left CI and Right CI are the confidence interval calculated by a one-LOD decrease from the estimated QTL position.

| Chromosome | Cluster | Position | LeftMarker | RightMarker | TraitName | LOD | PVE | Add | LeftCI |
| --- | --- | --- | --- | --- | --- | --- | --- | --- | --- |
| 2 | Tpa2.1 | 117 | solcap_snp_sl_8510 | solcap_snp_sl_15574 | plant width | 3.3851 | 6.4842 | 0.129 | 114.5 |
| 2 | Tpa2.1 | 117 | solcap_snp_sl_8510 | solcap_snp_sl_15574 | X mass | 4.1424 | 6.8279 | 0.139 | 114.5 |
| 3 | Tpa3.1 | 30 | solcap_snp_sl_9689 | solcap_snp_sl_9681 | plant area | 3.6327 | 8.0934 | 0.1522 | 24.5 |
| 3 | Tpa3.2 | 58 | SL3.0ch03-61078160 | solcap_snp_sl_10372 | plant width | 3.2538 | 9.4831 | 0.1555 | 56.5 |
| 3 | Tpa3.2 | 58 | SL3.0ch03-61078160 | solcap_snp_sl_10372 | X mass | 3.8291 | 9.4218 | 0.1628 | 56.5 |
| 3 | Tpa3.2 | 59 | SL3.0ch03-61078160 | solcap_snp_sl_10372 | CM height | 3.794 | 3.6957 | 0.1603 | 56.5 |
| 4 | Tpa4.1 | 128 | SL3.0ch04-1213025 | solcap_snp_sl_63869 | CM height | 3.8063 | 12.0286 | -0.2891 | 127.5 |
| 6 | Tpa6.1 | 10 | SL3.0ch06-13533085 | SL3.0ch06-3310762 | convex area | 4.741 | 11.2755 | 0.1621 | 8.5 |
| 6 | Tpa6.1 | 10 | SL3.0ch06-13533085 | SL3.0ch06-3310762 | plant width | 3.5865 | 6.9265 | 0.1336 | 8.5 |
| 6 | Tpa6.1 | 10 | SL3.0ch06-13533085 | SL3.0ch06-3310762 | plant area | 3.5117 | 7.3344 | 0.1457 | 8.5 |
| 6 | Tpa6.1 | 10 | SL3.0ch06-13533085 | SL3.0ch06-3310762 | X mass | 4.8686 | 8.016 | 0.151 | 8.5 |
| 6 | Tpa6.1 | 10 | SL3.0ch06-13533085 | SL3.0ch06-3310762 | Y mass | 4.902 | 9.3612 | 0.1757 | 8.5 |
| 8 | Tpa8.1 | 80 | solcap_snp_sl_21461 | SL3.0ch08-63373341 | X mass | 3.2439 | 5.3622 | 0.123 | 74.5 |
| 9 | Tpa9.1 | 36 | solcap_snp_sl_26683 | SL3.0ch09-4516570 | CM height | 4.8307 | 6.146 | 0.2081 | 32.5 |
| 9 | Tpa9.2 | 67 | SL3.0ch09-39293163 | SL3.0ch09-63338944 | convex area | 3.1976 | 7.523 | -0.1323 | 60.5 |
| 10 | Tpa10.1 | 71 | solcap_snp_sl_8835 | Le013158s_161 | Y mass | 3.2077 | 5.9734 | 0.1404 | 68.5 |

The parent that donated the favorable allele was determined according to the sign of the QTL additive effect (Awata et al., 2020), where a positive sign referred to H7996 resistant parent and a negative sign to WV susceptible parent. Typically, in QTL analysis, the favorable allele has the higher trait value. Here, a higher trait value is favorable in all cases except the visual wilting score, in which a higher trait value was associated with susceptibility (e.g. 90% wilting is more susceptible than 20% wilting). For QTL detected using our visual wilting score, the susceptibility allele was contributed by the susceptible parent WV (Table 1), consistent with other tomato-*R. solanacearum* QTL studies based on visual assessment of wilting. For all QTL except those on chromosome 2 and 10, the favorable allele was contributed by Hawaii 7996. QTL clusters on chromosome 2 and 10, which contributed to plant height and width, were donated by the susceptible parent WV. Favorable allelic contribution from both resistant and susceptible parents is common in QTL studies for resistance (Maschietto et al., 2017; Awata et al., 2020).

Among the 20 individual *Bwr* QTL were 10 QTL clusters (Table 1). We use the term ‘QTL cluster’ to describe QTL for different traits that co-localize at the same left and right genetic marker. There are between one and six *Bwr* QTL within a given cluster, and each QTL within the cluster has different LOD scores and explains a different percentage of phenotypic variation. *Bwr* QTL for traits that are highly correlated with each other (Figure 2B) tended to cluster together. For example, a cluster of QTL on chromosome 3 (*Bwr3.1*; Table 1) contains six *Bwr* QTL, including three for area-related traits (convex width, and X mass and CM width), which are correlated with each other at r = 0.94 - 0.97 (Figure 2B). In another cluster (*Bwr3.2*) on this same chromosome, *Bwr* QTL for traits that describe plant area were detected together. This co- localization supports the robust nature of our analyses.

In other clusters, only one trait that described one aspect of the wilting phenotype was present. For example, despite several metrics that describe width, only convex width was identified as a QTL on chromosome 4 (*Bwr4.1*; LOD 3.18; PVE = 6%). This suggests that the image-based phenotyping captured genetic variation that is specific to each trait.

### Image-based non-destructive phenotyping identifies three types of QTL clusters

Among our *Bwr* QTL clusters, we identified three types: those found using both image-based and visual phenotyping (1 cluster, *Bwr3.2*), those found only through visual phenotyping (1 cluster, *Bwr6.1*), and those that were identified only through image-based phenotyping (8 clusters; Table 1 and Figure 3).

**Figure 3:**
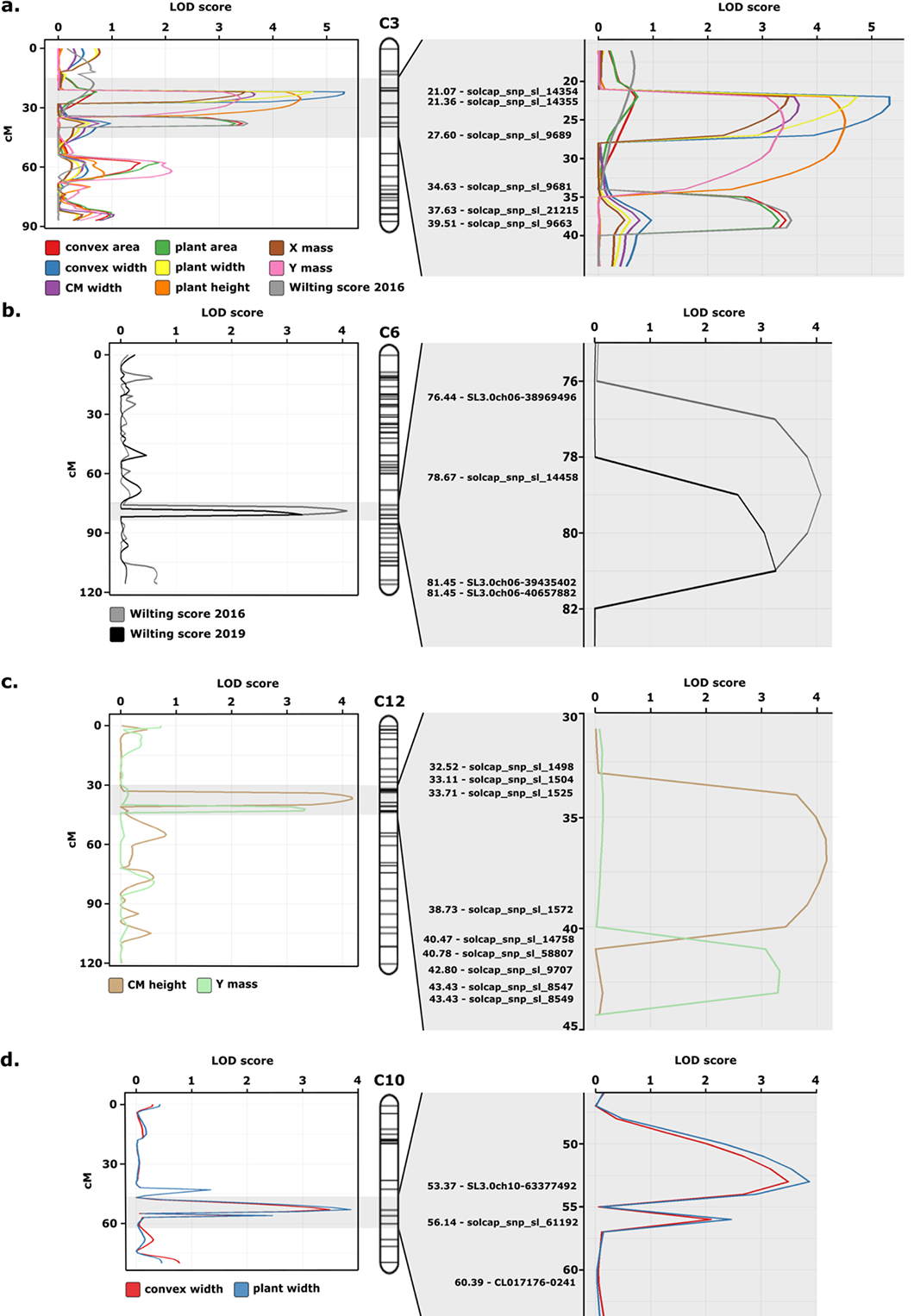
Significant QTL clusters on chromosomes 3 (a.), 6 (b.), 12 (c.) and 10 (d.). The image-based and visually assessed traits are represented with different colors for each chromosome. The vertical axis represents the genetic position (cM) and horizontal axis shows the LOD score. For each chromosome, the left panel represents the entire chromosome and the right panel represents the significant QTL cluster regions.

For the cluster including both image-based and visual phenotyping, (*Bwr3.2*) on chromosome 3, we detected QTL for convex area, plant area and visual plant wilting in 2016 (Table 1 and Figure 3a). Previous studies examining tomato responses to other strains of *Ralstonia* have not detected *Bwr* QTL in this region, indicating that it may be a novel target for *R. solanacearum* strain K60. The identification of both image-based and visual QTL within the same cluster suggests that our phenotyping methods are robust at identifying features of plant wilting. Close to this cluster is *Bwr3.1*, which detected QTL for traits based on plant width, including convex width and CM width. The proximal arm of chromosome 3 may be an important but unexploited region for defense against *R. solanacearum* strain K60.

On chromosome 6 we detected *Bwr* (*Bwr6.1*) using only visual assessment for both 2016 and 2019 (Table 1, Figure 3b). This region may be a ‘hot spot’ for QTL for resistance to multiple strains of *Ralstonia*. Previous studies using this same tomato population for mapping *Bwr* to other strains of *Ralstonia* detected QTL approximately 4 Mb from this position (Supplemental Figure 7).

In all other clusters, QTL were identified using only image-based phenotyping. Two such clusters, detected on chromosome 12, identified QTL for CM height (*Bwr12.1*) and Y mass (*Bwr12.2*), and were located near previous QTL detected against other strains of *Ralstonia* (Supplemental Figure 7 and Figure 3c) . These previous studies (Wang et al., 2000; Jaw-Fen Wang et al., 2013; Shin et al., 2020) detected QTL using visual wilt assessment of adult plants in the field.

The remaining QTL detected through image-based phenotyping have not been previously identified in studies using other strains of *Ralstonia*. These regions may be effective only against specific strains of *Ralstonia* (‘strain-specific’ QTL), or may include those that are not easily detected using visual assessment. In support of the latter, at one of the image-based-only clusters, *Bwr10.1* (Figure 3d), we detected a QTL using visual assessment that did not meet our threshold for significance after permutation (Supplemental Table 3). Thus, image-based phenotyping improved our ability to detect QTL.

### Image-based phenotyping identified Bwr QTL prior to visual symptom development

In our system, wilting symptoms begin to appear on highly susceptible plants at 4 dpi, and these plants are nearly 100% wilted by 6 dpi. To test whether we could detect *Bwr* QTL prior to the onset of visual symptoms, we performed QTL analysis at 3, 4 and 5 dpi. No QTL were identified based on visual assessment at any of these time points, however several QTL were identified based on image-based phenotyping. At 3 dpi, three clusters were detected on chromosome 3, two of which co-localized with those identified at 6 dpi (Supplemental Table 4). *Bwr3.1*, a cluster detected only by image-based phenotyping, and *Bwr3.2*, a cluster detected by both visual and image-based phenotyping, were first detected at 3 dpi. *Bwr3.1* was also detected at 5 and 6 dpi, while *Bwr3.2* was detected 3, 4, 5 and 6 dpi (Supplemental Table 4). A small number of additional QTL were detected at earlier time points (Supplemental Table 4), but in most cases these were not stable, since they were not present at 6 dpi. These data suggest that image-based phenotyping can identify *Bwr* QTL prior to the onset of wilting symptoms.

### Plant architecture QTL do not overlap with Bwr QTL

The parents of the RIL population, H7996 and WV, have different aboveground phenotypes (Supplemental Figure 4) and the shoot architecture of the RILs correspondingly varies. To ensure that *Bwr* QTL were the result of tomato responses to *Ralstonia*, and not due to differences in aboveground plant architecture, we used our traits in a QTL analysis at -1 dpi, the day before plants had been inoculated. We call these ‘tomato plant architecture (*Tpa*)’ QTL. We identified 16 *Tpa*, within 9 QTL clusters (Table 2). None of the 16 *Tpa* were detected within the same interval as *Bwr* at 6 dpi. However, two *Tpa* QTL clusters were detected that were near *Bwr* QTL present at 6 dpi. One of these was detected on chromosome 3 at position 30, between *Bwr3.1* and *3.2*.

**Table 2.** Overview of tomato plant architecture (Tpa) QTL identified at -1 dpi by image-based phenotyping.

| Chromosome | Cluster | Position | LeftMarker | RightMarker | TraitName | LOD | PVE | Add | LeftCI | RightCI |
| --- | --- | --- | --- | --- | --- | --- | --- | --- | --- | --- |
| 2 | Bwr2.1 | 45 | solcap_snp_sl_18519 | SL3.0ch02-49265134 | plant width | 3.4108 | 6.4576 | -0.1669 | 42.5 | 47.5 |
| 2 | Bwr2.2 | 72 | SL3.0ch02-45885016 | SL3.0ch02-44375028 | X mass | 3.4765 | 9.4924 | -0.2178 | 69.5 | 73.5 |
| 2 | Bwr2.2 | 72 | SL3.0ch02-45885016 | SL3.0ch02-44375028 | CM width | 3.2455 | 8.455 | -0.2189 | 69.5 | 73.5 |
| 3 | Bwr3.1 | 22 | solcap_snp_sl_14355 | solcap_snp_sl_9689 | convex width | 5.3238 | 10.2506 | 0.2103 | 21.5 | 26.5 |
| 3 | Bwr3.1 | 22 | solcap_snp_sl_14355 | solcap_snp_sl_9689 | plant width | 4.7309 | 9.1511 | 0.1976 | 21.5 | 25.5 |
| 3 | Bwr3.1 | 22 | solcap_snp_sl_14355 | solcap_snp_sl_9689 | X mass | 3.4765 | 5.8012 | 0.1705 | 21.5 | 26.5 |
| 3 | Bwr3.1 | 23 | solcap_snp_sl_14355 | solcap_snp_sl_9689 | CM width | 3.6627 | 6.2716 | 0.1887 | 21.5 | 27.5 |
| 3 | Bwr3.1 | 25 | solcap_snp_sl_14355 | solcap_snp_sl_9689 | plant height | 4.5107 | 7.6902 | 0.1896 | 21.5 | 31.5 |
| 3 | Bwr3.1 | 25 | solcap_snp_sl_14355 | solcap_snp_sl_9689 | Y mass | 3.3918 | 9.3283 | 0.1708 | 21.5 | 31.5 |
| 3 | Bwr3.2 | 38 | solcap_snp_sl_21215 | solcap_snp_sl_9663 | Wilting Score 2016 | 3.5258 | 7.3736 | -0.1726 | 35.5 | 39.5 |
| 3 | Bwr3.2 | 38 | solcap_snp_sl_21215 | solcap_snp_sl_9663 | convex area | 3.4328 | 9.0796 | 0.1953 | 34.5 | 39.5 |
| 3 | Bwr3.2 | 38 | solcap_snp_sl_21215 | solcap_snp_sl_9663 | plant area | 3.3019 | 8.6697 | 0.1902 | 34.5 | 39.5 |
| 4 | Bwr4.1 | 4 | SL3.0ch04-60431371 | solcap_snp_sl_11543 | convex width | 3.1855 | 6.0199 | 0.1627 | 1.5 | 10.5 |
| 5 | Bwr5.1 | 100 | solcap_snp_sl_22649 | SL3.0ch05-65105663 | plant height | 3.6115 | 9.4852 | 0.2103 | 95.5 | 103.5 |
| 6 | Bwr6.1 | 79 | solcap_snp_sl_14458 | SL3.0ch06-39435402 | Wilting Score 2016 | 4.077 | 8.7626 | -0.1871 | 76.5 | 81.5 |
| 6 | Bwr6.1 | 81 | solcap_snp_sl_14458 | SL3.0ch06-39435402 | Wilting Score 2019 | 3.2673 | 11.0473 | -0.1621 | 78.5 | 81.5 |
| 10 | Bwr10.1 | 53 | solcap_snp_sl_33168 | SL3.0ch10-63377492 | plant width | 3.8661 | 7.6897 | -0.1811 | 50.5 | 54.5 |
| 10 | Bwr10.1 | 53 | solcap_snp_sl_33168 | SL3.0ch10-63377492 | convex width | 3.4883 | 6.8155 | -0.1714 | 50.5 | 54.5 |
| 12 | Bwr12.1 | 37 | solcap_snp_sl_1525 | solcap_snp_sl_1572 | CM height | 4.1755 | 10.7222 | 0.1837 | 33.5 | 40.5 |
| 12 | Bwr12.2 | 42 | solcap_snp_sl_58807 | solcap_snp_sl_9707 | Y mass | 3.3249 | 8.5939 | 0.1651 | 40.5 | 43.5 |

While we cannot rule out the possibility that the same genes underlie these *Tpa* and *Bwr* QTL, their different marker intervals, coupled with the visual wilting QTL that is within the *Bwr3.2* cluster, suggests that different genes are responsible for these QTL. Together, these data suggest that our image-based phenotyping detected genomic regions that function in responding to *Ralstonia* and *Bwr* QTL are not the result of differential growth patterns within the RILs.

## Discussion

Breeding for plant disease resistance is one of the best strategies to combat plant diseases and prevent major crop loss, but is challenging in part due to the complicated nature of disease phenotyping. Here we used rapid, non-destructive, image-based phenotyping with RGB images to identify 20 QTL in 10 clusters for tomato responses to *Ralstonia* at 6 dpi, two of which were detected as early as 3 dpi. Together, the 20 *Bwr* QTL at 6 dpi explained more than 88% of the variation in response to *Ralstonia*. Image-based phenotyping for shape-based traits that were correlated with wilting detected both novel loci and those that overlapped with QTL based on human visual assessment. These results establish the importance and feasibility of quantitative, non-destructive, imaged-based phenotyping to identify new genetic targets for crop disease resistance during disease progression.

### Benefits of image-based phenotyping

Imaged-based phenotyping has been used to detect QTL associated with plant root (Topp et al., 2013) and shoot (Zhang et al., 2017; Knoch et al., 2020; Li et al., 2020) architecture, plant height (Wang et al., 2019), salt stress (Awlia et al.), and yield (Tanger et al., 2017; Pauli et al., 2016) among other traits. Although image-based phenotyping has become increasingly common to quantify plant disease symptoms (Mahlein, 2016; Mahlein et al., 2017, 2019; Lowe et al., 2017; Shakoor et al., 2017; Mochida et al., 2019; Mir et al., 2019; Pérez-Bueno et al., 2019; Pineda et al., 2021; Simko et al., 2017), few studies have used this technology to identify new genetic loci for plant disease resistance (Yates et al., 2019; Fordyce et al., 2018; Corwin et al., 2016). One reason for this may be that many plant disease symptoms occur at the leaf scale (such as spots and specks), making it difficult to use some sensors non-destructively and in high-throughput, or at the proper resolution needed to assess disease. For example, automated digital phenotyping of Septoria Tritici Blotch on wheat leaves identified novel QTL for resistance, but the destructive phenotyping required significant manual labor to harvest, mount and scan leaves (Yates et al., 2019). While wilting does occur at the leaf scale, it is an easier phenotype to assess at the whole plant scale. Another possibility is that imaging sensors are often expensive, making it more challenging to phenotype the large number of plants needed in a QTL or GWA study. Our method is low-cost and rapid ( < 2 min/plant), and requires no manual labor other than placing the plant on the turntable and initiating imaging via a computer.

Our image-based phenotyping identified loci that function in tomato responses to *Ralstonia*, but did not detect one of the two *Bwr* clusters (*Bwr6.1*) we found using visual assessment of disease. Visual assessment was performed at 8 dpi, while our image based phenotyping ended at 6 dpi. It is possible *Bwr6.1* is not apparent until very late stages of disease. Alternatively, these results could mean that we missed components of wilting in our image-based traits. However, a more likely explanation is that our image-based traits were averaged across eight two dimensional images, but wilting is a three dimensional (3D) phenotype. Images in 3D may allow detection of *Bwr6.1*. It would also be of interest to test whether using additional types of sensors, such as hyperspectral, would identify *Bwr 6.1*. Multi and hyperspectral imaging has been used to detect biochemical changes in disease (Mahlein et al., 2017; Lowe et al., 2017; Zhang et al., 2020). Given the chemical changes that occur during water stress, this type of imaging would likely identify additional genetic variation associated with resistance to bacterial wilt disease.

### New QTL for responses to Ralstonia in tomato

As in tomato, resistance to *Ralstonia* in other Solanaceous crops is quantitative (Young and Danesh, 1994; Danesh et al., 1994; Thoquet et al., 1996a, 1996b; Mangin et al., 1999; Wang et al., 2000; Carmeille et al., 2006; Jaw-Fen Wang et al., 2013). Breeding for resistance QTL is thus the primary way forward to developing *Ralstonia*-resistant crops in the Solanaceae. This has not been easy, in part because of the diversity of *Ralstonia*. The *Ralstonia solanacearum* species complex was recently subdivided into three species (*R. solanacearum*, *R. pseudosolanacearum*, and *R. syzygii*), each prevalent in a different part of the world (Remenant et al., 2012; Safni et al.; Prior et al., 2016). Each species has multiple strains, with an overlapping, but distinct set of virulence proteins that promote disease (Landry et al., 2020). Varieties with effective resistance will likely have QTL that are effective against local strains (strain-specific QTL) as well as those effective against multiple strains (broad-spectrum QTL).

Most previously identified QTL have focused on resistance to *R. pseudosolanacearum*, and none of them used the strain of *R. solanacearum* as in this work (Danesh et al., 1994; Thoquet et al., 1996a, 1996b; Wang et al., 2000; Carmeille et al., 2006; Jaw-Fen Wang et al., 2013; Shin et al., 2020; Mangin et al., 1999). Using visual assessment of wilting and the same RIL population of tomato (H7996 x WV700) used here, one broad-spectrum QTL for resistance to multiple strains of *R. pseudosolanacearum,* and one strain of *R. solanacearum* (JT-516) had been previously identified on chromosome 6 (Danesh et al., 1994; Thoquet et al., 1996a, 1996b; Mangin et al., 1999; Wang et al., 2000; Carmeille et al., 2006; Jaw-Fen Wang et al., 2013; Shin et al., 2020). We also identified a QTL on chromosome 6, using visual assessment in both 2016 and 2019. In our study, *Bwr6.1* confers 8 – 11% of the variation, compared to 11.5 – 33% for *Bwr6a - Bwr6d* (Jaw-Fen Wang et al., 2013; Shin et al., 2020). The genes within the left and right intervals of *Bwr6.1* are particularly interesting because there are two groups of NBS-LRRs located between the left and right markers for *Bwr6.1* (solcap_snp_sl_14458 and the GBS-identified SNP at 39435402). One of these groups includes the *S. lycopersicum* homolog of Arabidopsis *RESISTANCE to PSEUDOMONAS SYRINGAE 4* (*RPS4*). Fine-mapping of *Bwr6.1* is needed to determine whether a specific NBS-LRR protein or receptor is responsible for the resistance conferred by this region.

Previous studies also identified a major QTL on chromosome 12 that is effective for resistance to *R. pseudosolanacearum*, and explained between 15.9 – 53.9% of the variation (Shin et al., 2020). We found two clusters of QTL on chromosome 12, for image-based traits. Within the region spanning *Bwr12.1* on chromosome 12 is one NBS-LRR disease resistance gene (*Solyc12g10740*) and one receptor-like kinase (*Solyc12g010660*) while within *Bwr12.2* are genes related to disease resistance pathways, including a gene encoding a Pathogenesis Response (PR) Protein (*Solyc12g014310*), and two genes encoding Leucine Rich Repeat – Receptor Like Kinases (*Solyc12g014350* and *Solyc12g150105*). Our results suggest that this region of chromosome 12 may be important for resistance to multiple strains of *Ralstonia*. The QTL we identified on chromosomes 2, 3, 4, 5, 8 and 10 were not previously identified, and thus may be specific to *R. solanacearum* strain K60. With the exception of *Bwr3.2*, these QTL were detected using only our image-based traits.

Together, our results establish the value of image-based, non-destructive disease phenotyping for uncovering novel genetic components and new targets for quantitative disease resistance in crops. By identifying new genetic loci, this type of rapid phenotyping may enable the identification of broad-spectrum and durable resistance.

## Methods

### Plant growth

Seeds of 188 Hawaii7996 (H7996) x West Virginia700 (WV) recombinant inbred lines (RILs) in the F8 generation were obtained from the Asian Vegetable Research and Development Center (AVRDC) in June 2014. Seeds were propagated to the F9 generation in field and greenhouse in West Lafayette, IN in 2014 and 2015 and were used in QTL analysis in 2016 and 2019.

For all plants used in the imaging experiments and 2019 visual assessment of wilting, seeds were sown into individually labeled 1801 traditional inserts that were placed into 1020 flats (Hummert International, USA). Seeds in the 2016 experiment were sown into individually labeled 1203 inserts that were placed into 1020 flats. Seeds were sown into ProMix Propagation Mix supplied by the Lilly Greenhouses and Growth Facility located at Purdue University, West Lafayette, Indiana, USA. The individual pots were randomized and placed in a growth chamber at 28°C, relative humidity of 65% for a lighting cycle of 16 hours light/8 hours dark. The plants began germination on day 4 and plants were inoculated with *R. solanacearum* strain K60 at 17 days after planting when three true leaves were present. Trays were rotated in the growth chamber throughout each experiment. In 2016, four seeds of each RIL were grown in the growth chamber with parental controls, and each plant was visually assessed for wilting at 8 days post inoculation (dpi). In 2019, one seed per RIL, along with parental controls, was grown in the growth chamber for each of five independent replicates. Each plant was imaged at -1, 3, 4, 5, and 6 dpi, and visually assessed at 8 dpi.

For plants used to genotyping with tomato SolCap markers, F9 generation RIL seeds and parental controls were grown in the greenhouse in 2-gallon pots with Metro Mix 510 soil. Plants were grown for 6 weeks.

### Ralstonia solanacearum growth and plant inoculation

*Ralstonia solanacearum* strain K60 (containing a GFP reporter) was grown on casamino acid-peptone-glucose (CPG) agar containing tetrazolium chloride (TZC) in the dark for 48 hours at 28°C as in (Caldwell et al., 2017). Briefly, bacteria were resuspended in sterile water to a concentration of approximately 2 x 10^8^ colony forming units (CFU)/mL for each experiment. For each experiment, the concentration of inoculum was confirmed through dilution plating. Pots of three-leaf plants were lightly compressed to induce wounding similar to transplant handling in field conditions. 60 mL of inoculum was applied to the surrounding soil using a serological pipet.

### Visual assessment of bacterial wilt disease

Wilt scores were visually assessed 8 dpi. Wilt scores were calculated by counting the number of wilted true leaves divided by the total number of true leaves on the plant. Plants were scored with a 95% if the plant had all of its leaves wilted, excluding the topmost leaf. Plants were marked as 100% when the topmost portion of the stem was collapsed and wilted (Supplemental Figure 2).

### Plant imaging

Plants were imaged the day before inoculation (-1 dpi) and then imaged on 3, 4, 5, and 6 dpi. A Linco Linstor 2000-watt photo studio (Amazon, USA) was used as the backdrop, and Flora X fluorescent lighting (Amazon, USA) was used to create a small photo studio. Individual plants were placed on a Photocapture 360 turntable (Ortery Technologies) that was programmed to capture eight images around the plant (every 45°). The images were captured using an EOS 6D Canon camera with an EF 50mm f/1.4 USM lens on a stationary tripod. An imaging carriage was created with a square petri dish and a fiducial marker attached to each side. A total of five independent replicates for each RIL and the parental controls were imaged.

### Phenotyping Pipeline and Image Analysis

From the plant images, ten traits were acquired for QTL analysis. A detailed description of the acquisition process can be found in (Yang et al., 2021, 2020). All traits were extracted based on plant color and shape. To eliminate any color discrepancy between images, images were automatically color corrected using a fiducial marker consisting of a colored checkerboard with known physical dimensions and colors.

After color correction, plant pixels were segmented from the rest of the image by thresholding channels in the L*A*B* color space. The resulting segmentation mask was improved with image morphological operations to fill holes and remove noise generated by the thresholding. The stem of the plant was identified using two neural networks, Mask R-CNN and U-Net. By locating the stem of the plant, metrics were defined that reflect the inner morphology of the plant. To train the neural networks, stem segmentation ground truth data were generated using Adobe Photoshop and LabelMe (a Python-based annotation tool) to mark the location of the image pixels belonging to the stem of the plant. Further details about this procedure are described in (Yang et al., 2020, 2021).

The plant and stem masks were used in the subsequent color and shape analysis, which acquired ten traits: total area of the plant mask (plant area), height of the plant mask (plant height), maximum width of the plant mask (plant width), area of the convex hull (convex area), width of the convex hull (convex width), color-based weighted average (color average), horizontal distance between the center of mass of the left and right sides of the plant stem (CM width), height of the center of mass (CM height), x-axis distribution of the center of mass (X mass), Y-axis distribution of the center of mass (Y mass).

The plant area, plant height and plant width were calculated from the plant mask. When calculating plant height, the upper 5% of the plant material was not included, which helped eliminate the impact of small leaves growing at the top. Using functions available in *OpenCV* Python Library, a convex hull was fit around the plant mask and the convex area and convex width were calculated. For color analysis, the weighted average of the pixel values in the A* channel was calculated, as this channel captures the green-magenta spectrum.

To calculate traits involving the center of mass (CM), the plant mask was split into left and right halves using the stem mask (Yang et al., 2020), and the CM located for each half. The x-axis distance between CMs (CM width) and the average height of the CMs (CM height) were used as traits.

The extension of just one leaf can have a major impact on measured plant width, generating a disproportionately larger value. Capturing the plant material distribution inside the plant mask can overcome this issue. Using the split plant mask, the horizontal and vertical distribution of plant material was estimated. The distances at which 90% of the plant material was captured in the horizontal and vertical directions were used as the traits ‘X mass distribution’ and ‘Y mass distribution’. The average of eight views around the plant were used for each trait value. Outputs are in pixels.

In the random forest classification, visual wilting scores were thresholded to 0 or 1, where 0 = 0 to 50% wilting, 1 = 51 – 100% wilting. Average trait values from eight views from 6 dpi for 969 plants (RILs and parental controls) were used across all five replicates. Some plants did not have a trait value for that day and were not used in the algorithm.

### DNA extraction, Marker generation and Genotyping by Sequencing (GBS)

Tomato SolCap markers and single nucleotide polymorphisms (SNPs) identified through GBS were used for map creation. Leaf disc samples were collected from each RIL plant using a biopsy punch and were sent to LGC, Biosearch Technologies. Genomic DNA was extracted and genotyped for 128 SNPs from the Tomato SolCAP panel using LGC’s KASP assay. One hundred twelve SNPs were obtained for RILs using this method.

For GBS, genomic DNA (gDNA) was extracted from 188 F9 individuals of the RIL population and from parental plants using Trizol. gDNA was RNase-treated and cleaned with phenol chloroform extraction. Sequencing library preparation was as described by (Elshire et al., 2011). Briefly, gDNA was digested with *PstI* and 150 bp paired end sequencing was performed on two lanes of an Illumina Hi-seq 2500 at the Purdue Genomics Facility.

Reads were mapped to the *Solanum lycopersicum* 3.0 genome using the methods of (Manching et al., 2017). In brief, reads were de-multiplexed, and adapter sequences were removed. Reads were filtered based on the presence of a GBS barcode on the forward read (R1), which accounts for 95.4% of the PE reads. For these R1, 97.3% had a paired R2 read, and 2.7% did not have a pair. R1 read pairs and R1 singles reads were combined and assessed independent of pairing and treated as single end reads for the remaining analyses (924,500,737 reads).

Reads were filtered to remove those with the presence of an internal restriction site, the lack of restriction site hang sequence at the end, and for minimum length, which retained 93.0% of the reads. There was a minimum of 17,561 reads, a maximum of 16,127,292 reads, and an average of 4,480,149 reads per sample (Supplemental Figure 8). Reads were mapped using BWA-MEM for paired end reads, and the GATK haplotype caller was used to generate a genomic Variant Call Format (gVCF) file for downstream analysis.

There were 74,082 SNPs called between the population and *Solanum lycopersicum* 3.0 reference genome. Many of these were SNPs between the reference genome and the population and do not vary in the population used here. Filtering for the presence of two alleles within the RIL population identified resulted in 2,738 SNPs. Of these, the parental alleles were identified in both parents for 278 SNPs and in one parent for 698 SNPs. The remaining 1,762 SNPs could not be assigned a parental origin and were not used for further analysis. The 976 SNPs with parental origin identified were filtered for a minor allele frequency greater than 0.02 and less than 0.99, which resulted in 632 high quality SNPs.

High quality SNPs identified from GBS were combined with previously defined SolCap markers from LGC Biosearch Technologies for a total of 748 markers. SNP coverage of the 188 RIL taxa was high. Of the 748 markers, the minimum coverage was 115 markers, the maximum coverage was 555 markers, and the average was 366 markers (Supplemental Figure 9). There was some residual heterozygosity in the taxa (Supplemental Figure 9b), and taxa with 10% or greater heterozygosity were removed from further analysis (2 RILs).

### Linkage Map Construction

The software QTL IciMapping (Meng et al., 2015) (version 4.1) was used for the map construction with all 748 markers. Redundant markers were filtered by taxa coverage using a missing rate of 85% and a distortion threshold at 0.001 to obtain a total of 408 markers. The 408 markers without anchor information were assigned to 12 groups based on a LOD score threshold value of 3. After grouping, the markers were ordered using the nearest neighbor algorithm (nnTwoOpt) using the rippling criterion SARF (Sum of Adjacent Recombination Frequencies) with a window size of five markers. After ordering, some markers at the end of the chromosomes were deleted when they were adding an insignificant genetic distance to the chromosome. These markers were identified after splitting the current chromosome in two sub-chromosomes between the longest marker interval. If the shortest sub-chromosome contained more than 20% of the markers before splitting, the two sub-chromosome were re-assembled. Otherwise, the shortest sub-chromosome was deleted.

### QTL detection

QTL were detected using Inclusive Composite Interval Mapping with additive effects (ICIM-ADD) in IciMapping (Meng et al., 2015) (version 4.1) using a genetic mapping with a 1 cM scanning step and a probability in stepwise regression of 0.001. The LOD significance threshold to declare a QTL significant was determined using a Type-I error of 0.0500 after one thousand permutation. For the QTL analysis, we removed 19 of the 188 RIL (10%) showing the highest variation according to the visual wilting score and 3 RIL due to missing data across the 5 biological repetitions for a total of 166 RIL into the QTL map (Supplemental Figure 3). The evolution of the ten image-based traits and two visual wilting scores between -1 dpi and 3, 4, 5 and 6 dpi were determined for the 166 RIL. To assure a normal distribution of the phenotypic data, the function “orderNorm” was used to perform an ordered quantile normalization (Peterson and Cavanaugh, 2020) before QTL analysis using the package “bestNormalize” version 1.6.1 with R software version 3.6.1 (R Core Team).

## Acknowledgements

We thank Chloe Caldwell for help with imaging, and Kristina Gans for help with GBS.

This work was funded by a Purdue Seed Grant to AIP and ED, a Foundation for Food and Agriculture Research New Innovator Award to AIP, and the endowment of the Charles William Harrison Distinguished Professorship at Purdue University to ED.

## Author Contributions

A.I.P. and E.D. designed the research; V.M., D.C., and B.K. performed research; V.M, E.E., S.B., C.Y, A.I.P, and E.J.D. analyzed data; S.B., C.Y., and E.D. contributed new computational tools; A.I.P. and V.M. wrote the manuscript; all authors edited and approved the manuscript.

---

**Supplemental Figure 1:**
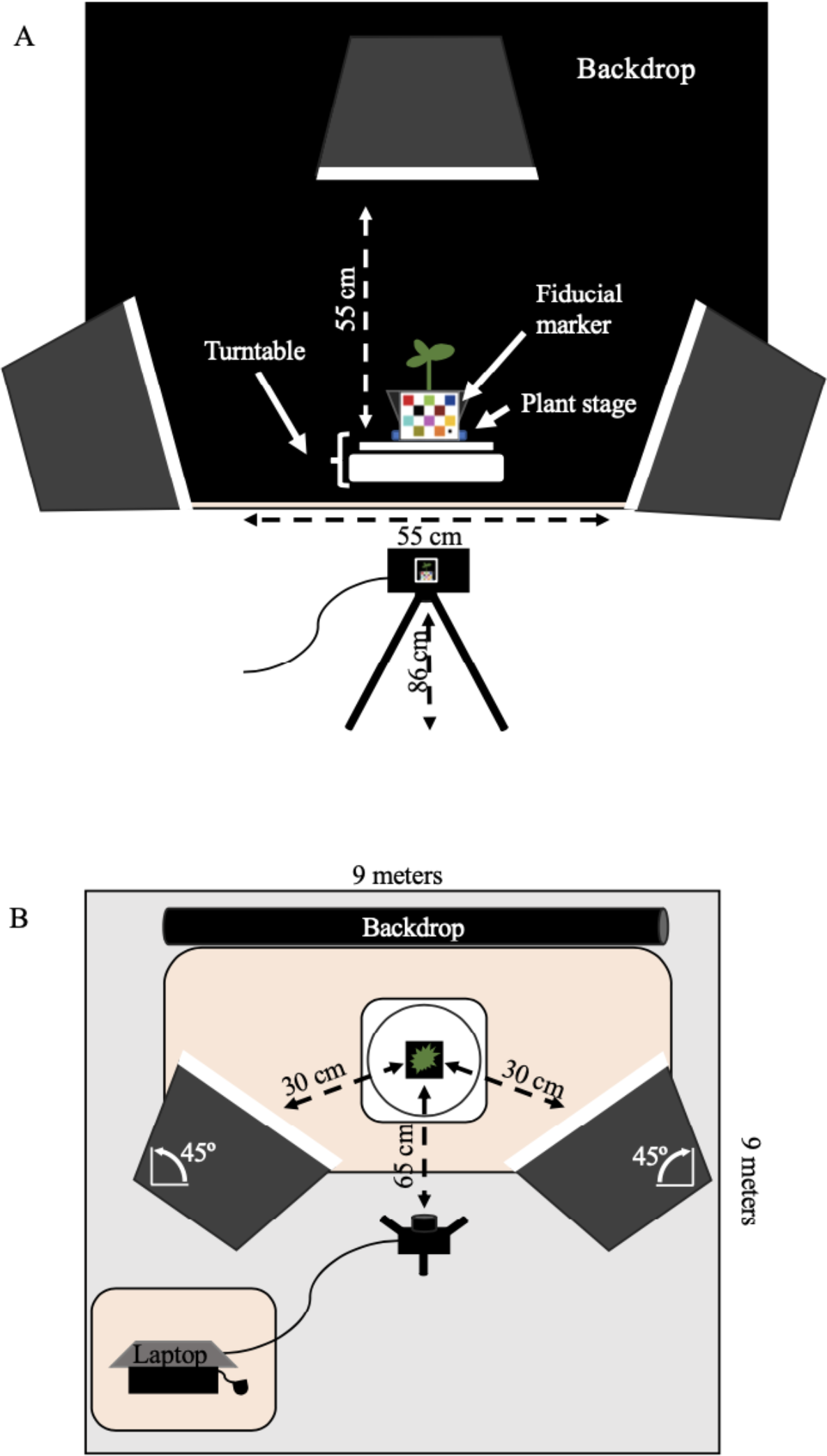
Design of our low cost phenotyping platform including automatic turntable, backdrop, lightning and RGB camera.

**Supplemental Figure 2:**
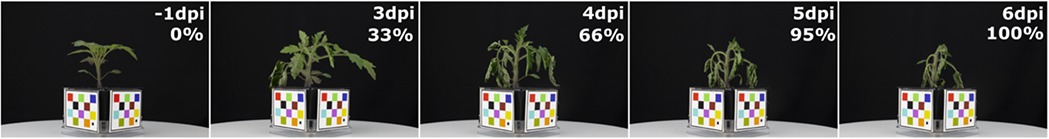
Raw RGB pictures showing the evolution of wilting symptoms on RIL # 646 at -1, 3, 4, 5 and 6 dpi. Visually assessed wilting score are expressed in percentage of wilted leaves.

**Supplemental Figure 3:**
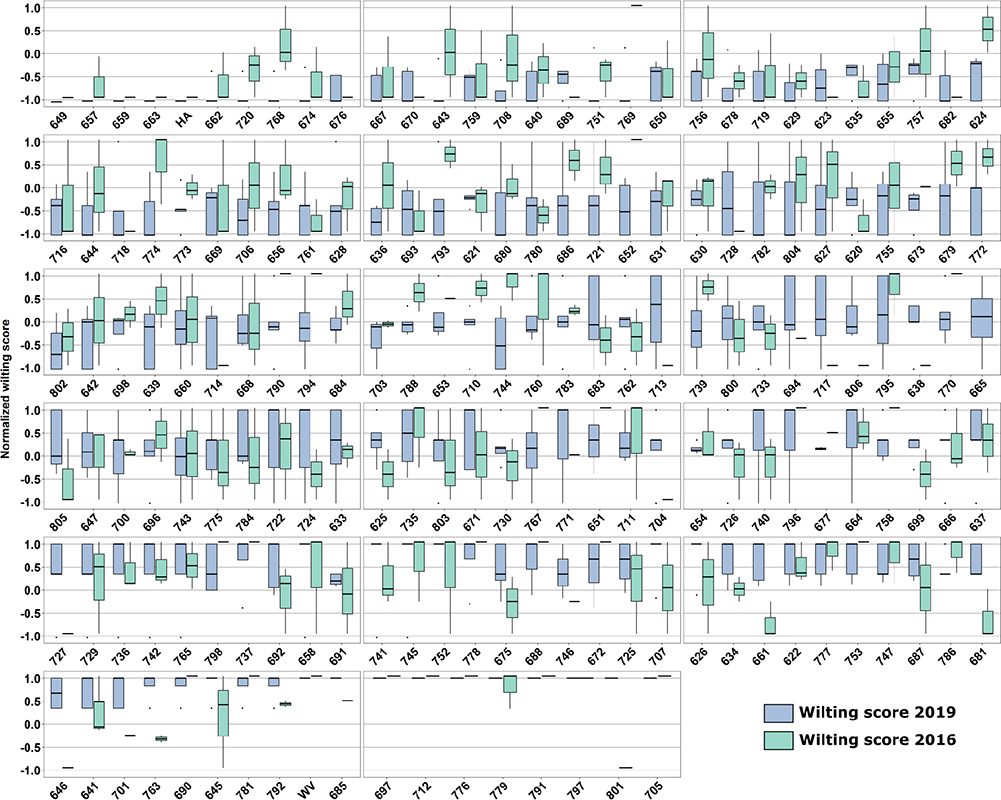
Wilting Score RIL: Boxplots showing the normalized wilting score at 6 dpi for the 166 RILs, resistant Hawaii 7996 (HA) and susceptible West Virginia (WV) genotypes.

**Supplemental Figure 4:**
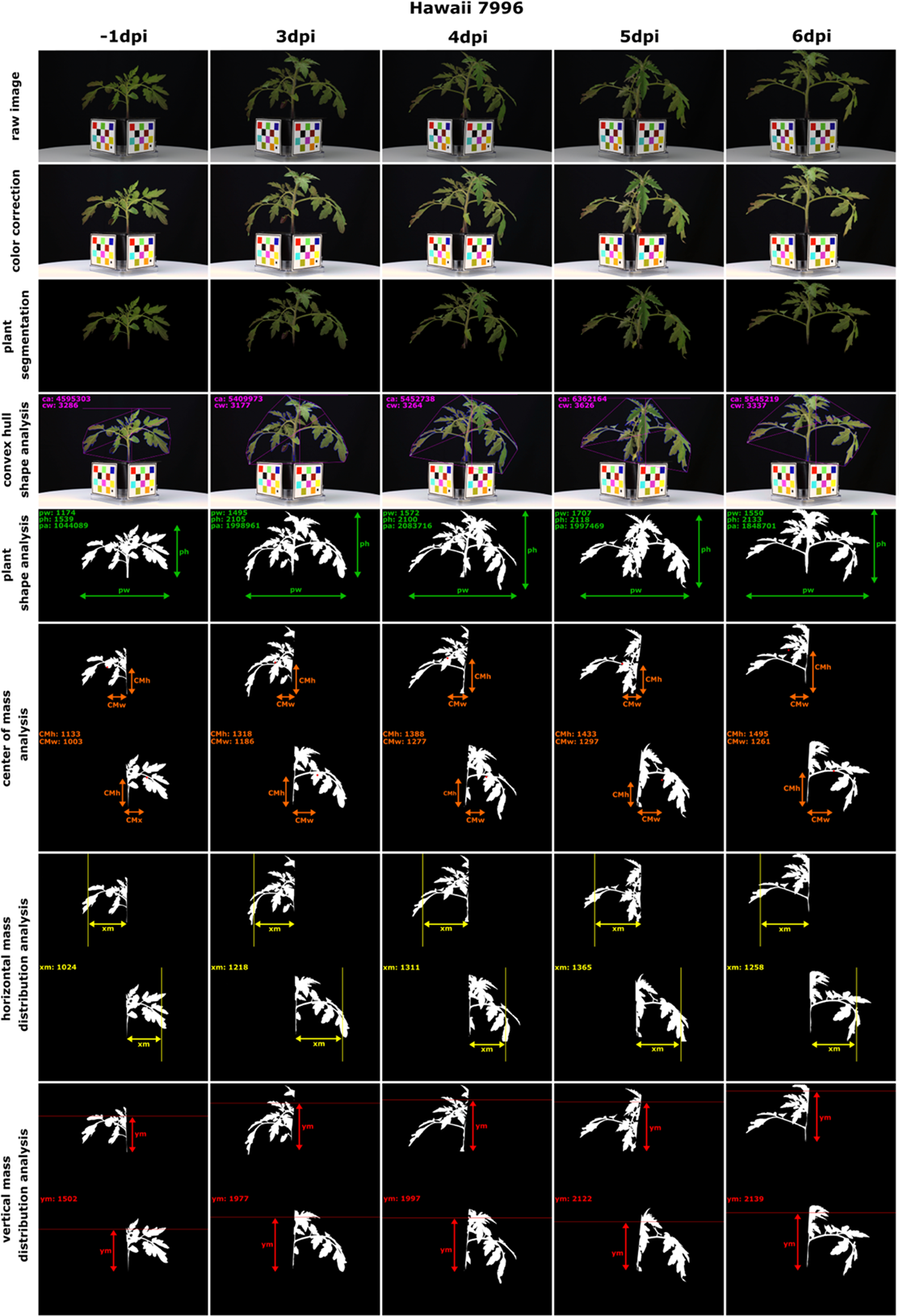
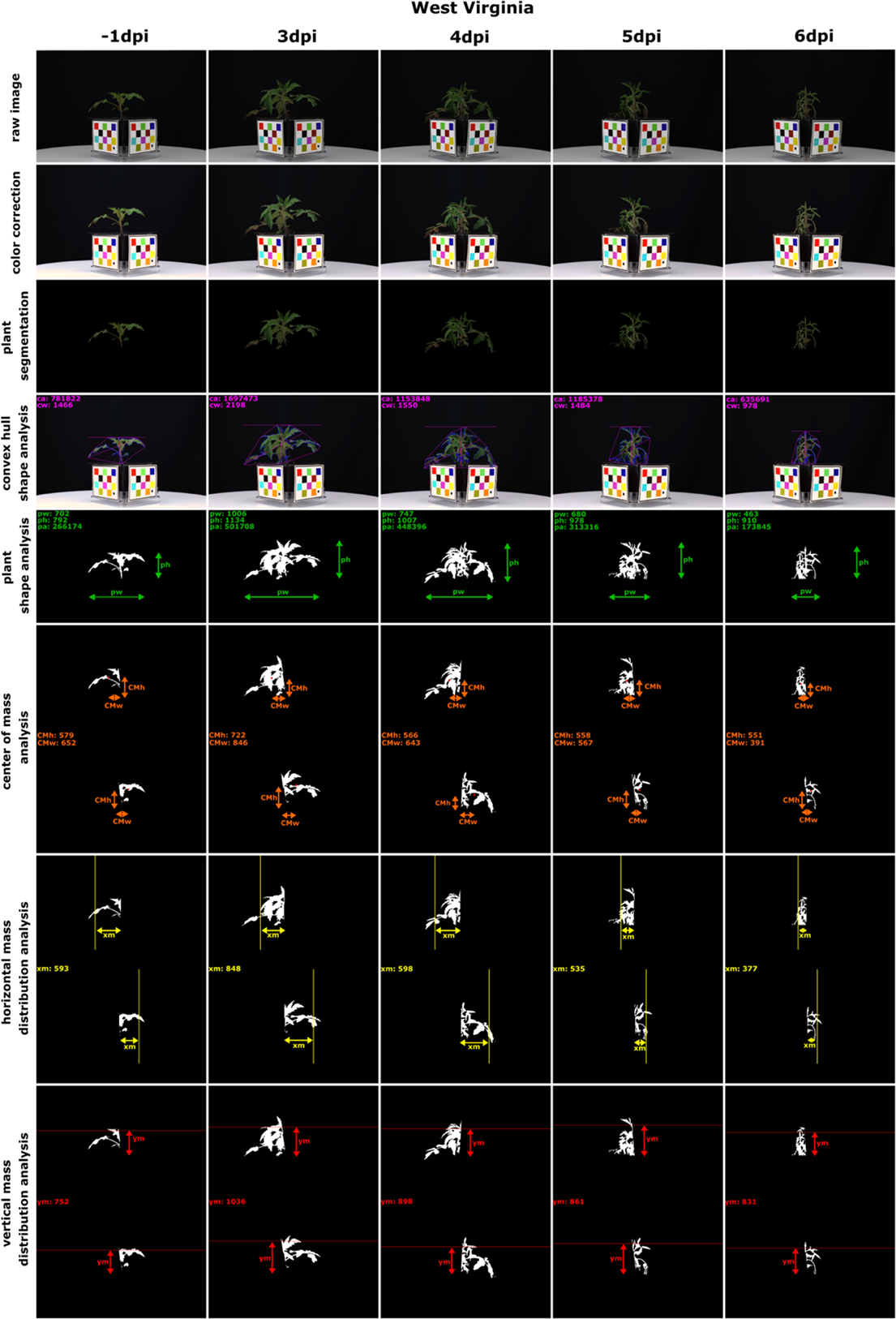
RGB pictures showing the evolution of the traits at -1, 3, 4, 5 and 6 dpi for A) Hawaii 7996 and B) West Virginia (pw: plant width, ph: plant height, CMh: Center of Mass height, CMw: Center of Mass width, xm: Horizontal mass distribution, ym: Vertical mass distribution). The Hawaii 7996 plant shown here is the same plant used for Figure 1.

**Supplemental Figure 5:**
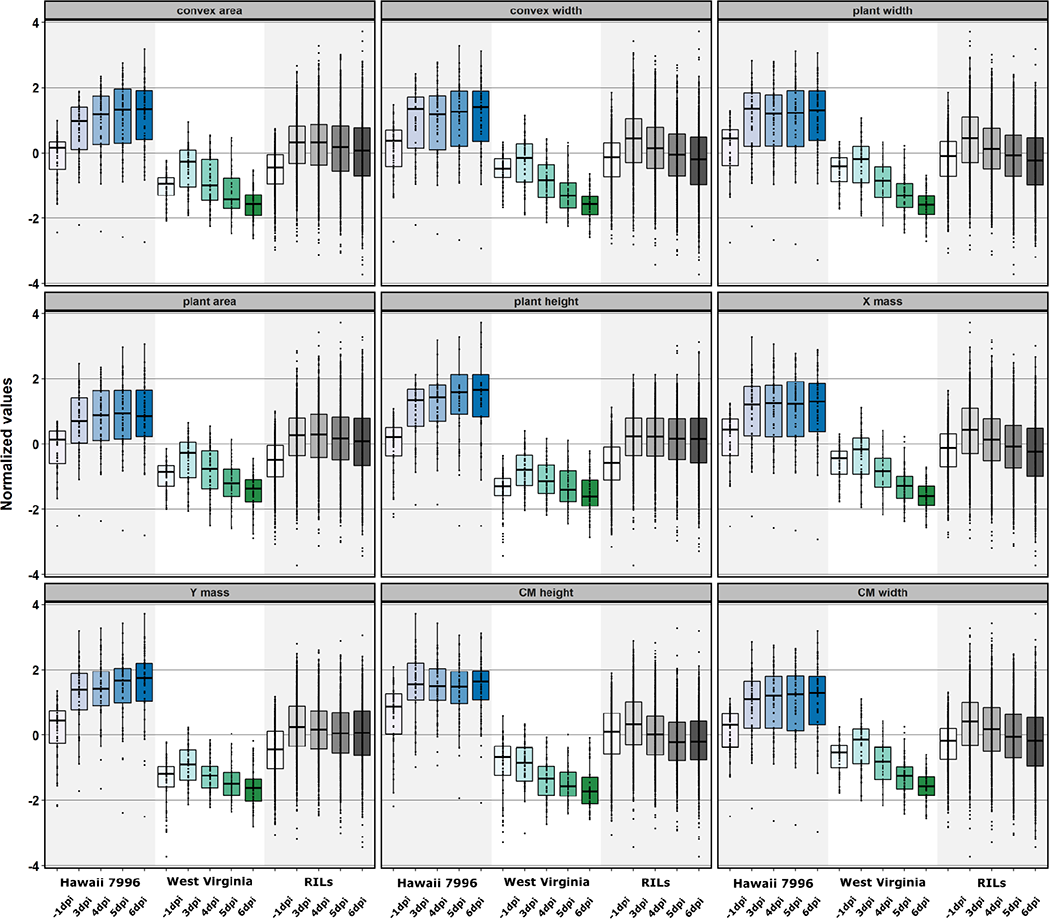
Boxplots showing the normalized values for image-based traits at -1, 3, 4, 5, 6 dpi for resistant Hawaii 7996, susceptible West Virginia 700, and 166 individuals of the RIL population.

**Supplemental Figure 6:**
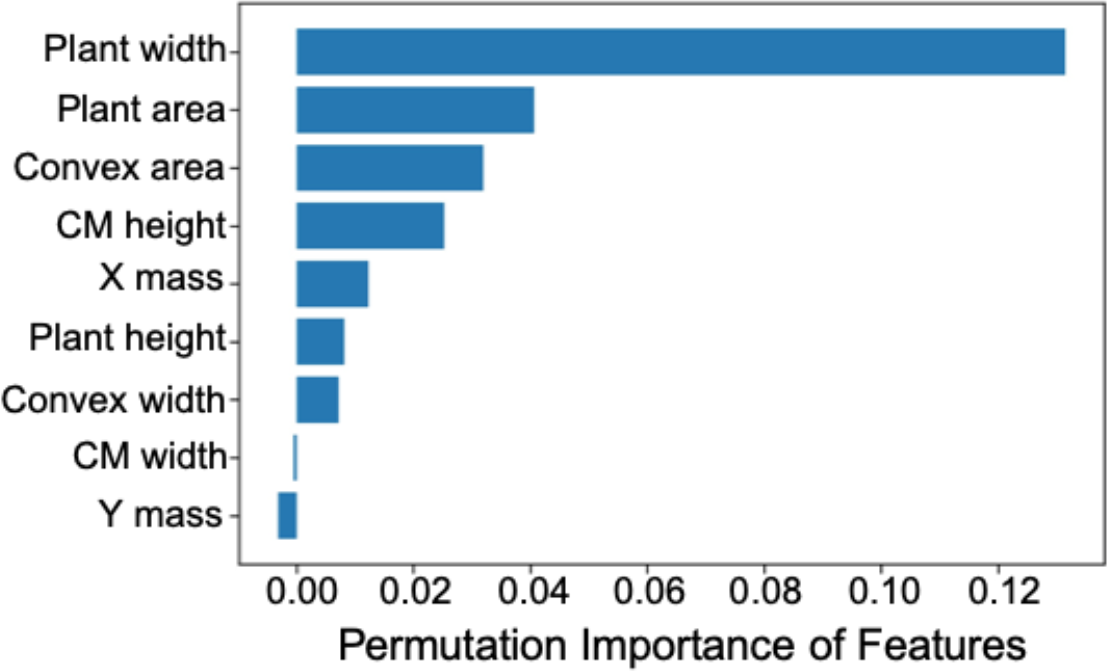
Histogram showing the image-based descriptors which contribute the most to wilting

**Supplemental Figure 7:**
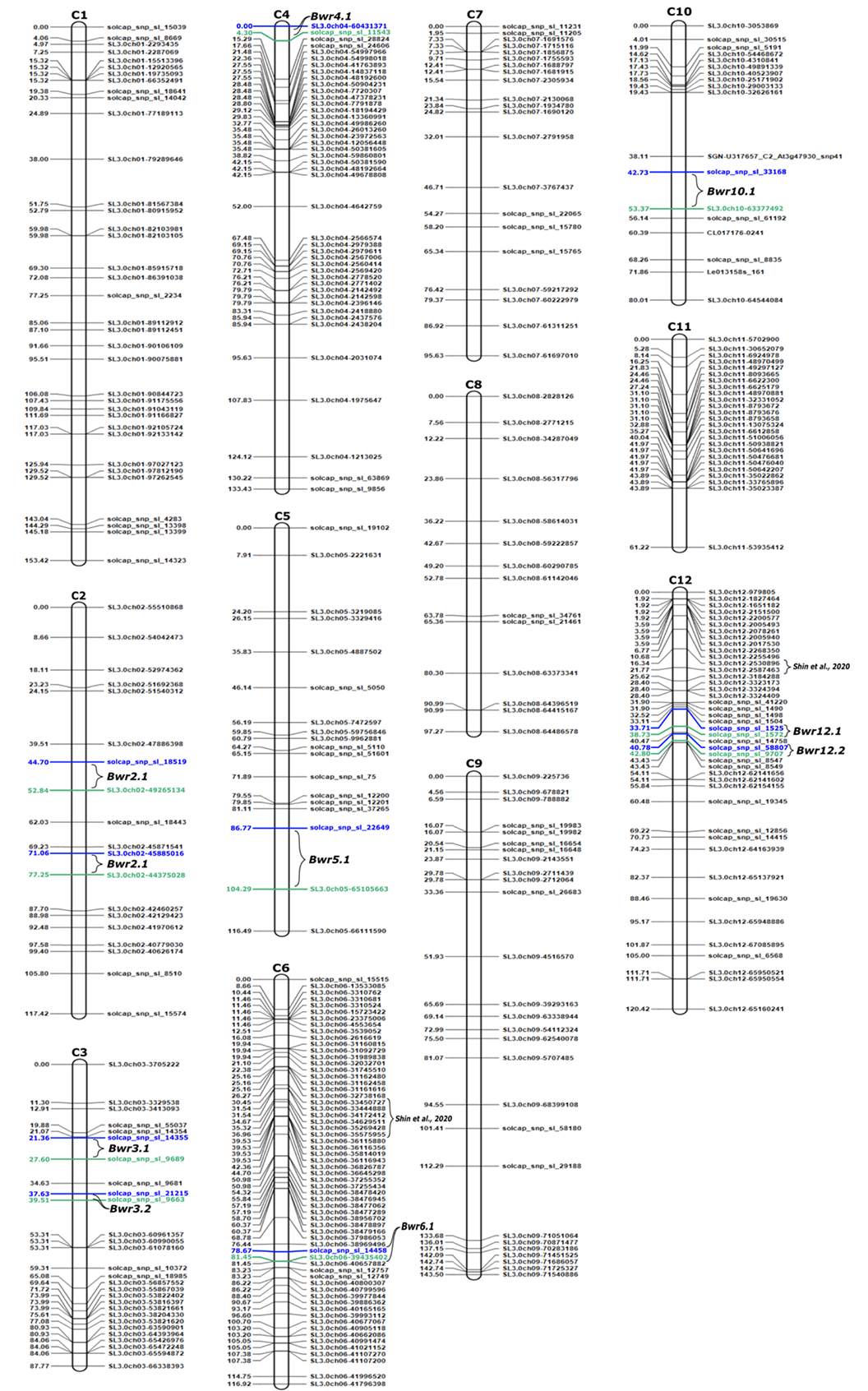
Genetic Linkage map constructed with SolCap and SNP markers. Location of 10 QTL clusters are displayed using blue and green for left and right markers respectively. Location of previously identified QTL on chromosome 6 and 12^27^ are also displayed.

**Supplemental Figure 8:**
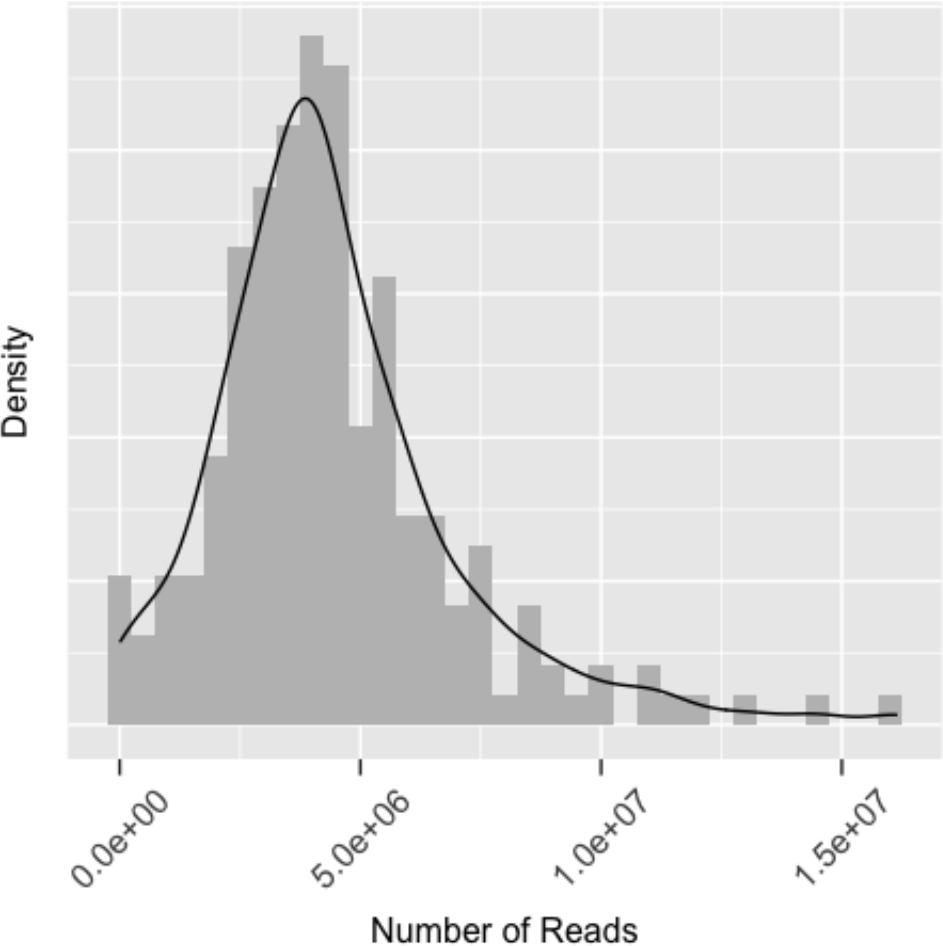
GBS Read Distribution by Sample (includes parental references).

**Supplemental Figure 9:**
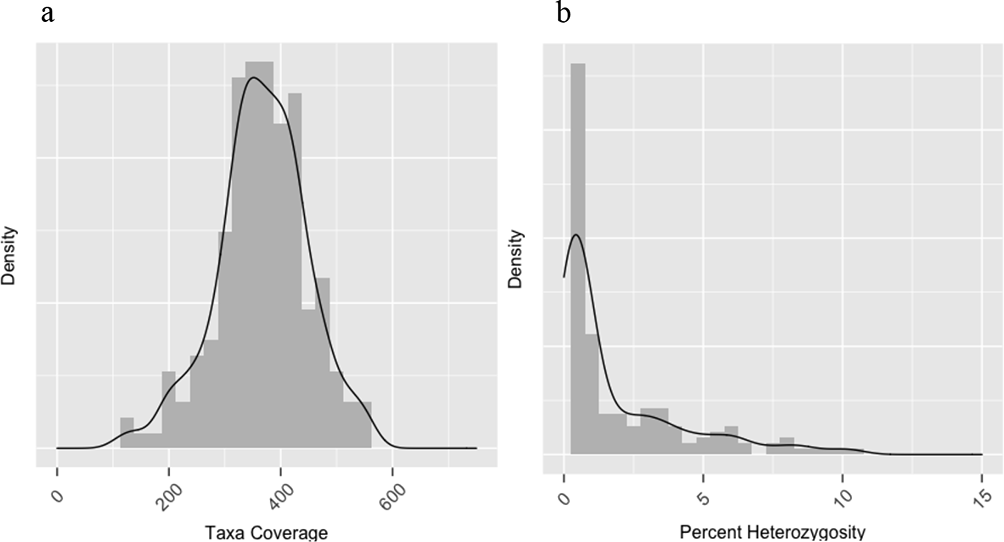
a) RIL and parental coverage and SNP density from GBS analysis; b) Percent heterozygosity and SNP density

**Supplemental Table 1.**
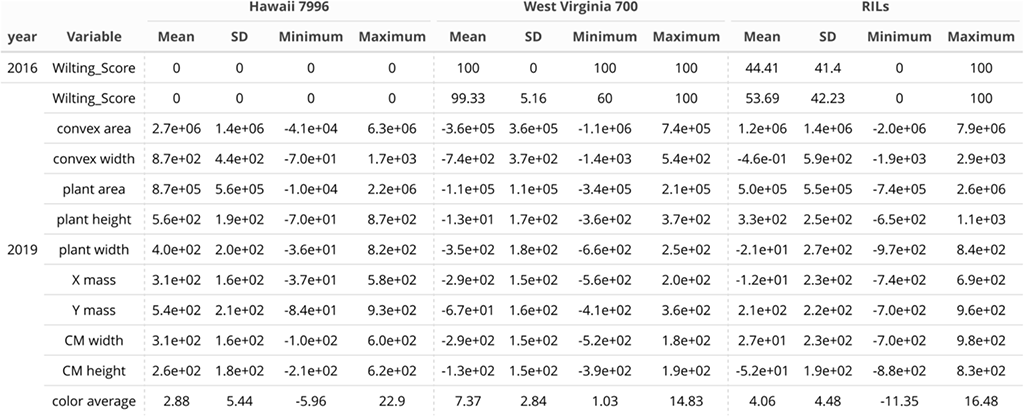
Descriptive statistics for 10 image-based traits and two years of visually assessed wilting scores (2016 and 2019). Image-based traits were from the same plants as the visually assessed wilting score in 2019. The RIL population was inoculated with Ralstonia and visually scored for wilting in 2016, but images were not taken. Data is shown for resistant parent Hawaii 7996, susceptible parent West Virginia 700, and the RIL population.

**Supplemental Table 2:** Marker density per cM for each chromosome.

| Chromosome | Number of marker | Chromosome Length (cM) | Average interval length (cM) | Standard deviation | Standard error |
| --- | --- | --- | --- | --- | --- |
| 1 | 36 | 153.42 | 4.38 | 4.15 | 0.70 |
| 2 | 19 | 117.42 | 6.52 | 3.92 | 0.92 |
| 3 | 28 | 87.77 | 3.25 | 3.56 | 0.68 |
| 4 | 44 | 133.43 | 3.10 | 4.25 | 0.65 |
| 5 | 18 | 116.49 | 6.85 | 5.33 | 1.29 |
| 6 | 61 | 116.92 | 1.80 | 1.99 | 0.26 |
| 7 | 21 | 95.63 | 4.78 | 3.94 | 0.88 |
| 8 | 14 | 97.27 | 7.48 | 4.44 | 1.23 |
| 9 | 27 | 143.5 | 5.52 | 5.77 | 1.13 |
| 10 | 18 | 80.01 | 4.71 | 4.80 | 1.16 |
| 11 | 25 | 61.22 | 2.55 | 3.88 | 0.79 |
| 12 | 43 | 120.42 | 2.87 | 3.01 | 0.46 |

**Supplemental Table 3:** Overview of the 9 genomic regions identified with a LOD score greater than 3 on chromosome 10 after inoculation with R. solanacearum at 6 dpi. According to the ICI Mapping analysis using 1000 permutations, a threshold value of 3.48 represents the minimum LOD to identify a significant QTL. Genomic regions with a lower LOD value did not reach our threshold for significance after permutation.

| Chromosome | Position | LeftMarker | RightMarker | TraitName | LOD | PVE | Add |
| --- | --- | --- | --- | --- | --- | --- | --- |
| 10 | 51 | solcap_snp_sl_33168 | SL3.0ch10-63377492 | Wilting Score 2019 | 3.0067 | 13.3810 | 0.2077 |
| 10 | 52 | solcap_snp_sl_33168 | SL3.0ch10-63377492 | Wilting Score 2019 | 3.0927 | 10.5192 | 0.1842 |
| 10 | 53 | solcap_snp_sl_33168 | SL3.0ch10-63377492 | Wilting Score 2019 | 3.0832 | 8.0396 | 0.1611 |
| 10 | 52 | solcap_snp_sl_33168 | SL3.0ch10-63377492 | convex width | 3.1634 | 9.0040 | -0.1835 |
| 10 | 53 | solcap_snp_sl_33168 | SL3.0ch10-63377492 | convex width | 3.4883 | 7.8538 | -0.1714 |
| 10 | 51 | solcap_snp_sl_33168 | SL3.0ch10-63377492 | plant width | 3.0398 | 11.1872 | -0.2039 |
| 10 | 52 | solcap_snp_sl_33168 | SL3.0ch10-63377492 | plant width | 3.5411 | 10.2424 | -0.1951 |
| 10 | 53 | solcap_snp_sl_33168 | SL3.0ch10-63377492 | plant width | 3.8661 | 8.8147 | -0.1811 |
| 10 | 53 | solcap_snp_sl_33168 | SL3.0ch10-63377492 | X mass | 3.0584 | 7.0293 | -0.1629 |

**Supplemental Table 4:**
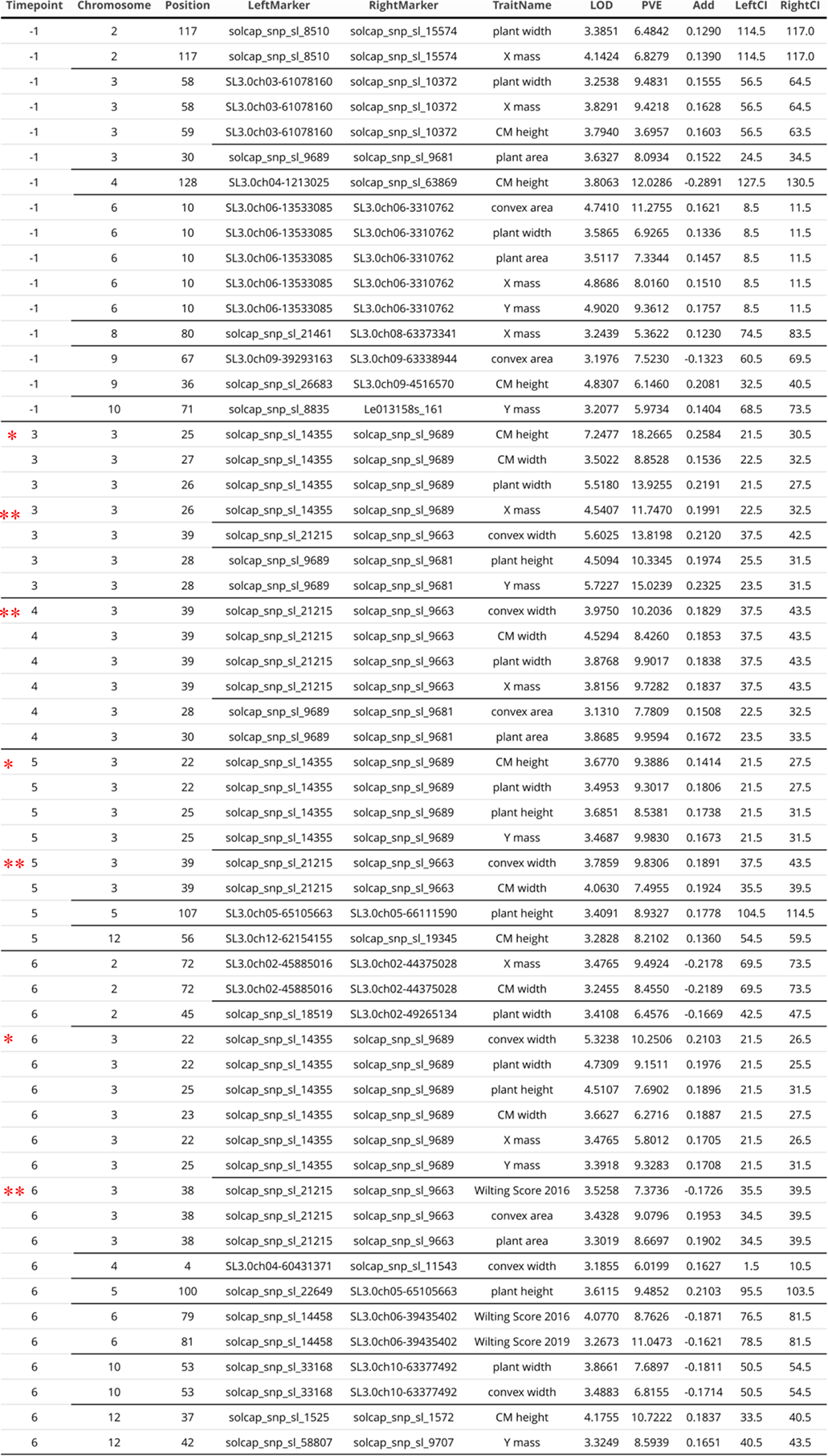
Overview of QTLs identified across the genome at all time points (-1, 3, 4, 5, 6 dpi) with R. solanacearum. LOD: maximum value of the Logarithm of the odd. PVE: Percentage of phenotypic variance explained. Add: Additive effect, the positive and negative values indicated that the alleles are introgressed from the resistant parent (Hawaii 7996) and susceptible parents (WestVirginia). LeftCI and RightCI are the confidence interval calculated by a one-LOD decrease from the estimated QTL position. * = Bwr3.1; ** = Bwr3.2

